# The post-hoc montage of perception: deep layers of primary visual cortex encode postdictive percepts

**DOI:** 10.64898/2026.09.04.749457

**Authors:** Pieter Barkema, Christoph Koenig, Joost Haarsma, Peter Kok

## Abstract

One of the most puzzling aspects of perception is postdiction: later information can change how a previous stimulus is perceived [1,2]. This indicates that perception is not a livestream of the external world, but rather a post-hoc reconstruction. How does the brain revise perception? It has been proposed that recurrent processing in the sensory hierarchy may sustain earlier information long enough for later information to impact it [3]. We hypothesised that this allows feedback from higher-order multisensory regions to alter the representation of earlier visual stimuli in the primary visual cortex (V1) [4]. Here, we tested this hypothesis using two postdictive illusions [5], where the presence or absence of a tone determines whether an illusory flash is perceived, or an actually presented flash is perceptually suppressed, respectively. This allowed us to isolate neural signals reflecting illusory percepts from sensory inputs in retinotopic space. We collected 7T layer-specific fMRI data while participants (N=23) experienced the illusions. We found that postdictive illusions induced similar activity patterns as real percepts, specifically in the deep layers of V1, suggesting postdictive feedback. Informational connectivity analyses suggested that the source of this feedback was the Superior Temporal Cortex (STC). We conclude that the postdictive montage of perception is reflected in V1, implemented through a delaying feedback loop with higher-order sensory cortex. This extends the role V1 plays in perceptual inference: beyond receiving sensory inputs reflecting the present and predictions of the future, it is involved in the post-hoc revision of perception.

## Introduction

Perception is the process whereby the brain tries to infer the current unknown state of the world based on a combination of sensory information and assumptions derived from prior experience [6]. Little is known, however, about how the brain integrates these two sources of information across time. The scientific dialogue has thus far mainly focused on how the brain integrates *previous* information into perception, i.e., *prediction* [7–9]. *Postdiction*, however, is a complementary phenomenon where information coming in *later* than the stimulus – within a timeframe of several hundred milliseconds – influences how it is perceived [3]. Postdiction shows us that – counterintuitively – our perception is not a livestream of the external world but is rather constructed in the brain post-hoc [10,11]. By studying the neural mechanisms of postdiction we can shed light on how the brain generates our conscious experience.

How is it possible that perception can be changed by later incoming information? There must be a delay at some stage of sensory processing that allows later information to catch up with earlier signals, creating a time window in which both sources can be integrated. To explain this delay, two plausible, yet untested, theories have been suggested [3,4]. The feedforward theory rests on the fact that the temporal integration window of neurons increases along the cortical hierarchy [12], allowing later information to catch up with earlier signals in higher-order sensory processing regions (e.g., in the temporal lobe). The feedback theory on the other hand proposes that recurrent feedback down the cortical hierarchy offers an opportunity for earlier signals to be influenced by later information in primary sensory cortex. That is, after the initial feedforward sweep, sensory signals are transmitted back to primary sensory cortex with a delay, due to reciprocal connections from higher sensory areas, allowing later information to modulate it. Distinguishing between these theories would provide insight into how our perceptual experience is constructed in the cortical hierarchy.

One crucial distinction between the two theories is the role of primary sensory cortex in postdictive perception. That is, the feedback theory predicts that primary sensory cortex would reflect postdictively revised perception, whereas the feedforward theory predicts that it would not, since integration would only take place in higher-order sensory regions with extended temporal receptive fields. Outside of the context of postdiction, it has been proposed that feedback to sensory cortex plays an important role in constructing perception [9,13,14]. For instance, retinotopically specific feedback to primary visual cortex (V1) has been shown to underlie the Kanizsa illusion [15–18]. We hypothesized that this crucial role of feedback to sensory cortex generalizes to postdictive perception.

To test this hypothesis, we employed two robust cross-modal postdictive illusions [5]. In one illusion, a percept of a visual flash is induced by an auditory stimulus in a retinotopic location that is not otherwise stimulated, and in the second illusion a presented flash is perceptually suppressed by the absence of an auditory stimulus (Figure 1A). Together, these illusions provide an ideal testbed to probe potential postdictive signals in V1, as they allow one to fully dissociate illusory percepts from retinal inputs (Figure S1). We hypothesized that the illusion would be reflected in the deep layers of V1, as the result of feedback from multisensory cortical areas (i.e., superior temporal cortex [4]) [15,19,20]. We tested this hypothesis by measuring layer-specific BOLD signals with ultra-high resolution 7T fMRI (0.8 mm voxels), combined with population receptive field (pRF) mapping [21] to isolate the region of V1 that was tuned to the location of the illusory flash.

**Figure 1.**
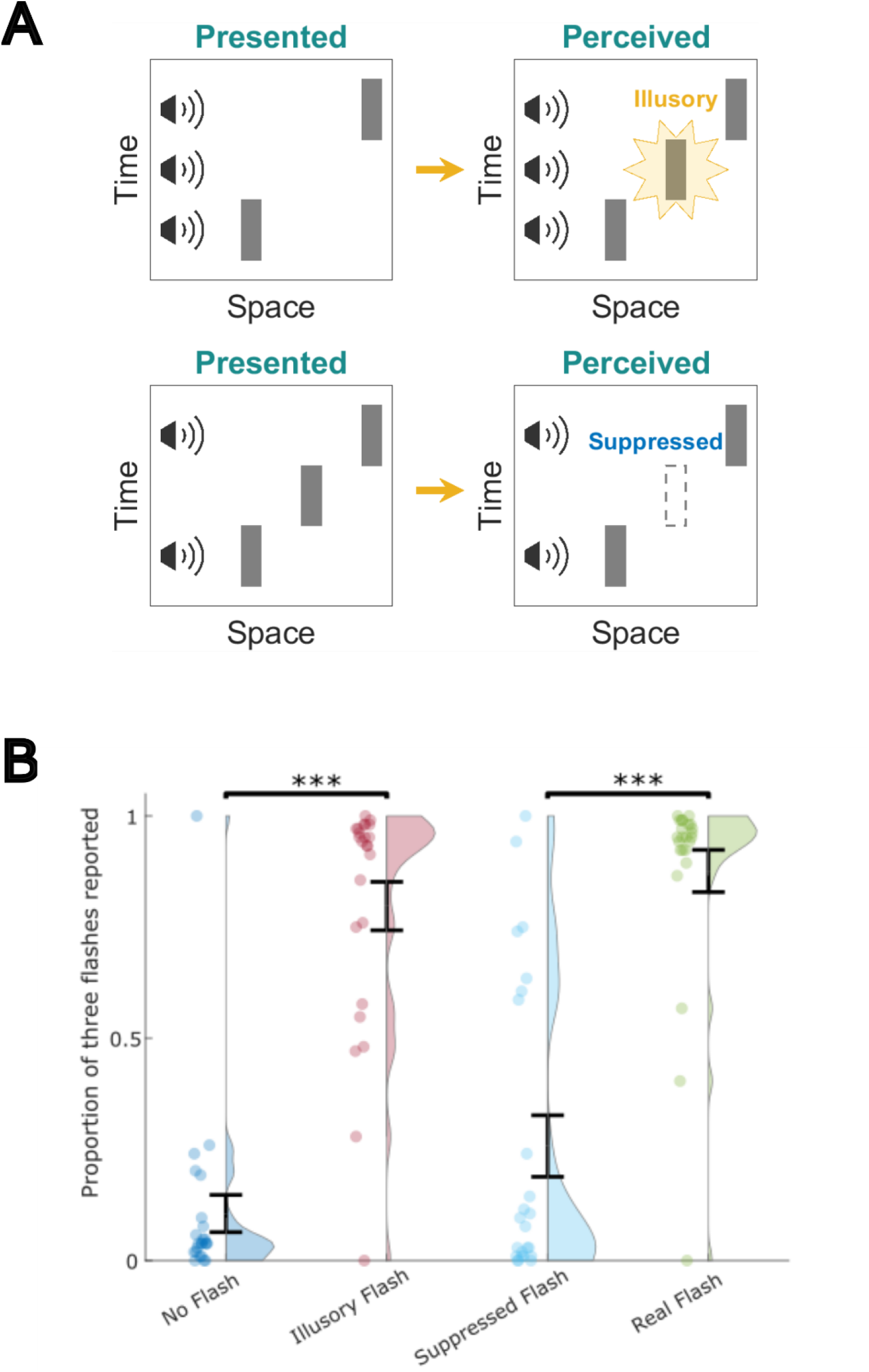
Experimental paradigm and behavioural results. A) Top: presenting two audiovisual stimuli pairs with an extra sound in between results in the perception an extra visual stimulus; bottom: presenting two audiovisual stimuli pairs with an extra flash in between result in suppression of this extra flash (adapted from Stiles et al. [5]). B) Proportion of trials on which participants reported seeing three vs. two flashes, separately for the different conditions. For a visual depiction of the conditions, see Fig S1. Black bars indicate SEM. *** *p* < 0.001.

To preview our findings, we found evidence that post-hoc constructed perceptual content was reflected in the deep layers of V1, in support of the feedback model of postdiction. These findings are in line with pre-perceptual sensory information in primary visual cortex being post-hoc corrected by perceptual inference [10,11].

## Results

We successfully induced two audio-to-visual postdictive illusions during fMRI scanning, and validated the control conditions (Figure 1B). In the ‘Illusory Flash’ condition, the middle sound induced an illusory flash, with three flashes being reported on .83 ± .04 (mean ± SEM) of trials despite only two being presented. This proportion was significantly higher than on trials without the extra sound (i.e., the ‘No Flash’ condition, .08 ± .02,; W(22) = 3.615, *p* < .001). In the second illusory condition, ‘Suppressed Flash,’ the absence of a sound suppressed the perception of a real flash, with three flashes reported on only .25 ± .07 of trials, significantly less than on trials with three flashes and three sounds (i.e., the ‘Real Flash’ condition, .92 ± .03; W(22) = 3.601, *p* < .001. Participants were highly confident in both their veridical and illusory percepts (Figure S2).

We asked whether these postdictive illusions were reflected in BOLD activity patterns in the layers of V1. For each postdictive illusion, we computed the ‘perceptual pattern similarity’, defined as the correlation between the BOLD activity pattern evoked by an illusion and the activity pattern evoked by the corresponding veridical percept (i.e., Illusory Flash x Real Flash and Suppressed Flash x No Flash), minus the correlation with the pattern evoked by the corresponding retinal input (i.e., Illusory Flash x No Flash and Suppressed Flash x Real Flash; see Figure 1A for visualization). This measure quantified whether the activity evoked by an illusion was more similar to perceiving the same thing (perceptual pattern similarity > 0) or to having the same input to the retina (perceptual pattern similarity < 0). We calculated this metric for both illusions, restricted to V1 voxels with a receptive field on the illusory location (i.e., the location of the middle flash).

This analysis revealed that the postdictive illusions affected the different layers of V1 layers distinctly (two-way mixed-effects ANOVA with factors ‘Layer’ and ‘Illusion Type’, main effect of Layer: F(1,21) = 4.588, *p* = .016; Figure 2A). Specifically, perceptual pattern similarity was stronger in deep layers than middle layers (t(21) = 2.735, *p* = .012). Other layer comparisons were not significant (deep vs. superficial: t(21) = 1.755, *p =* .093; middle vs. superficial, t(21) = 1.489, *p =* .151). In sum, only the deep layers showed a significant positive perceptual pattern similarity effect (one-sided t-test, t(21) = 2.027, *p =* .028; Figure 2), whereas this perceptual effect was not present in the other layers (mid: t(21) = −1.516, *p* = .144; superficial: t(21) = 0.414, *p* = .683), providing evidence that perceiving a postdictive illusion affects deep layer activity. This effect was not present in V1 voxels tuned to a control retinotopic location (all layers *p* > .05; Figure 2B). This striking effect demonstrates that the deep layers of V1 represented postdictively inferred percepts, rather than veridically reflecting retinal inputs.

**Figure 2.**
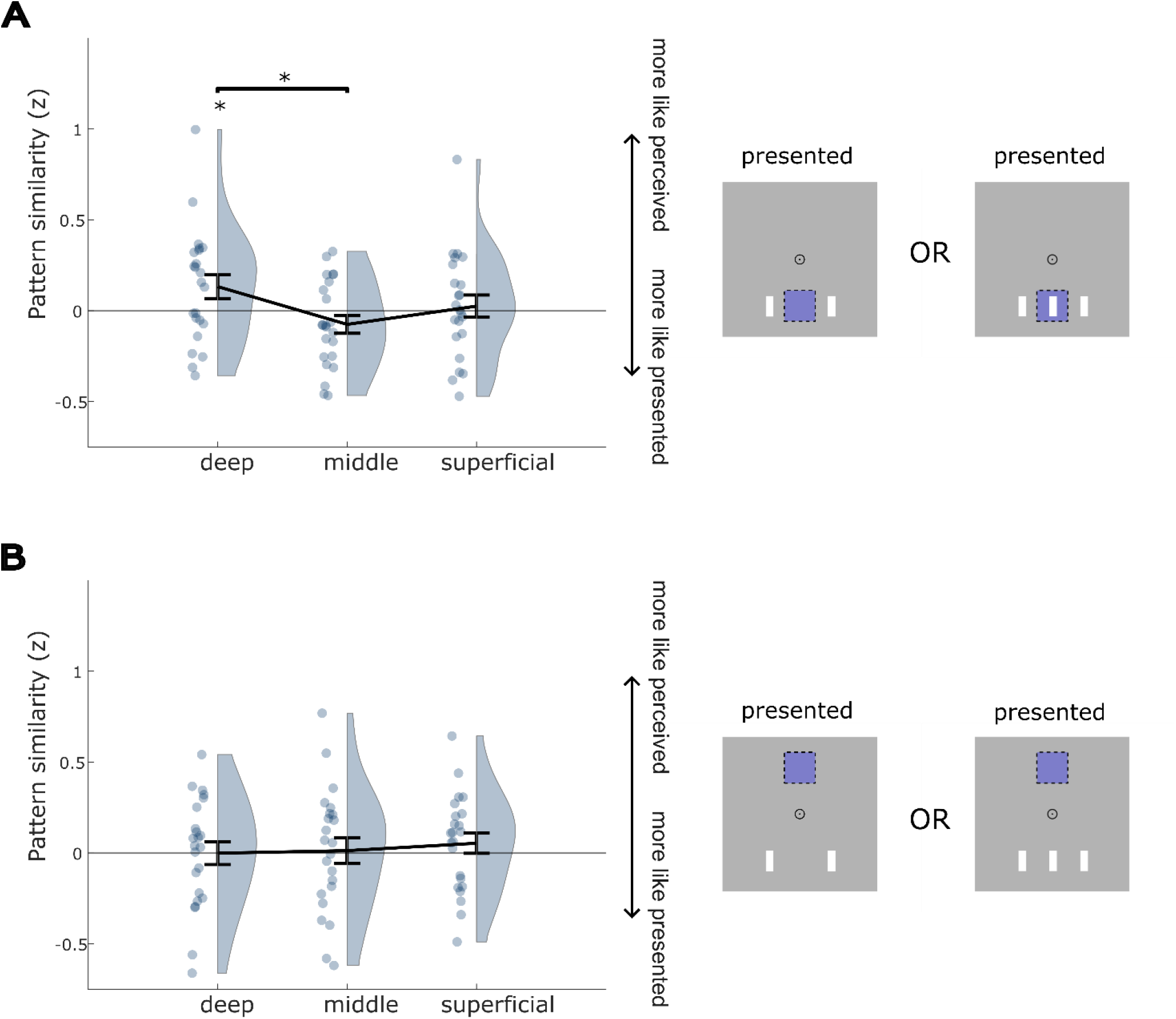
Perceptual pattern similarity in BOLD activity patterns for voxels tuned to the middle flash (A) and control (B) location per layer of V1, averaged over both postdictive illusions. Positive values indicate that activity patterns evoked by illusions were more similar to activity patterns evoked by stimuli inducing the same percept (e.g., Illusory Flash x Real Flash), while negative values indicate more similarity to conditions with identical retinal stimulus but different percepts (e.g., Illusory Flash x No Flash). Error bars denote SEM across participants. * *p* < 0.05.

The two illusions did not significantly differ in their effect on V1 activity (ANOVA, main effect of ‘Illusion type’: F(1,21) = 2.457, *p =* .132; interaction between ‘Layer’ and ‘Illusion type’: F(1, 21) = 1.999, *p* = .148), suggesting that the postdictive illusory creation and suppression of flashes, respectively, modulated V1 activity to a similar extent. That is, in both cases V1 deep layer activity more closely resembled the perceived stimulus than the retinally presented stimulus. To follow this up, we analysed the results for the two illusions separately. As expected, activity in V1 deep layers during the ‘Illusory Flash’ condition resembled activity evoked by a ‘Real Flash’ more strongly than activity evoked by ‘No Flash’ (one-sided t-test: t(22) = 1.814, *p* = .042; Figure 3A). This effect was stronger in the deep layers than the middle layers (t(22) = 3.644, *p* = .001). If anything, the middle layers showed a trend in the opposite direction (two-sided t-test: t(22) = −2.045, *p* = .053), in line with the fact that the middle layers receive feedforward sensory signals but not top-down feedback [22]. Activity patterns in the superficial layers did not clearly distinguish between perceptual and retinal similarity (t(22) = 0.108, *p* = .457), possibly since they contain a mixture feedforward and feedback signals (see Discussion). Importantly, these effects were specific to voxels with a receptive field on the illusory flash location, and were not present for voxels with a receptive field on a control region above fixation (all layers *p* > .05; Figure 3B).

**Figure 3.**
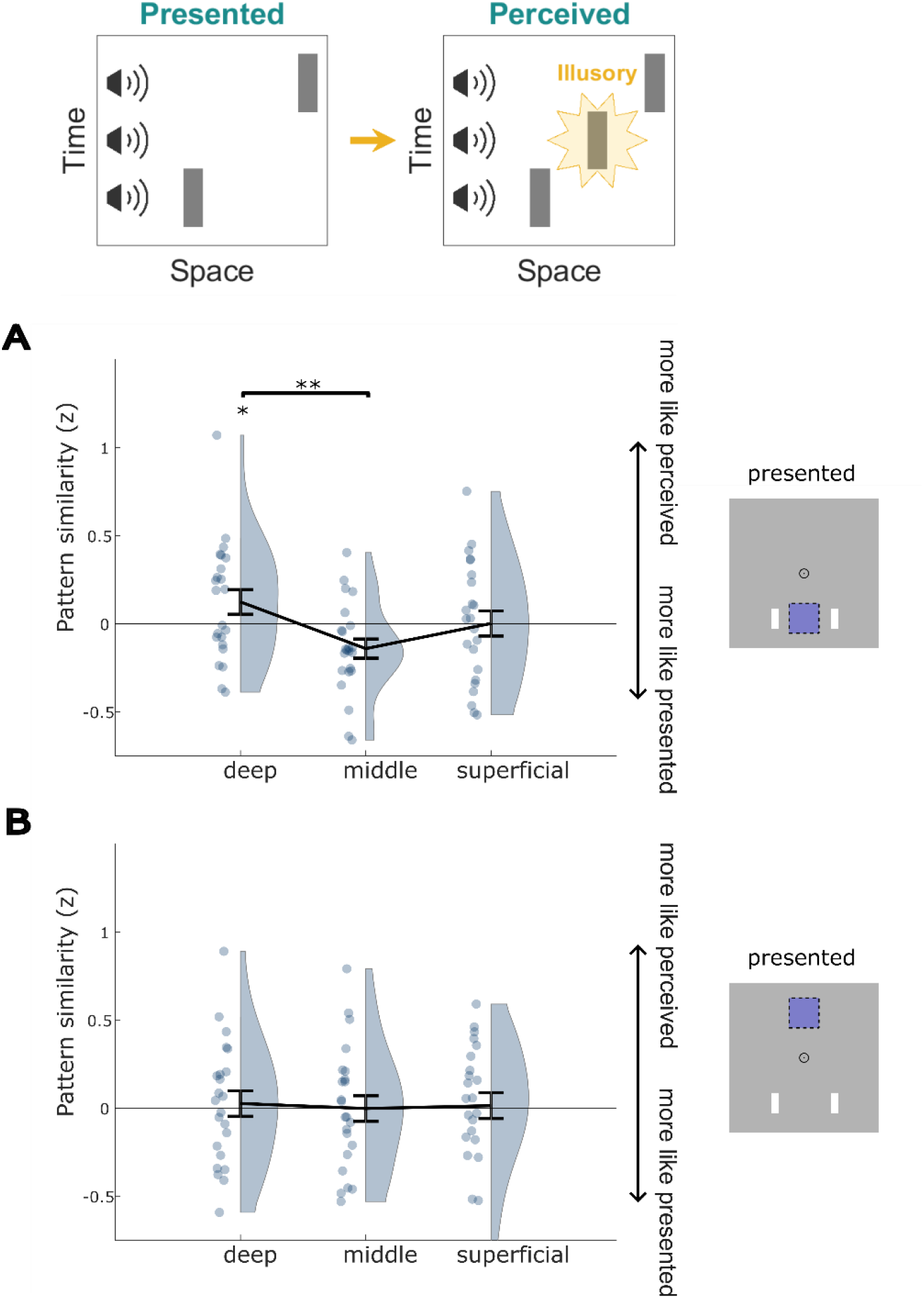
Perceptual pattern similarity in BOLD activity patterns for voxels tuned to the second flash and control location per layer of V1 for the ‘Illusory Flash’ illusion. A) Perceptual pattern similarity of voxels tuned to the illusory flash location. Positive values indicate that activity patterns were more similar to activity patterns evoked by stimuli inducing the same percept (Real Flash), while negative values indicate more similarity to conditions with identical retinal stimulus but different percepts (‘No Flash’). B) The same analysis for voxels tuned to a control retinotopic location. Error bars denote SEM across participants. * *p* < 0.05; ** *p* < 0.01.

We performed the same analysis on the ‘Suppressed Flash’ trials (Figure 4). It should be noted that this condition provides a less strong test of our hypotheses, for two reasons. First, the illusory absence of a percept occurs in the presence of retinal input, i.e., a strong bottom-up drive, and neural signals in V1 are thus a mixture of retinal inputs and perceptual absence, rather than a ‘pure’ inferential signal as in the Illusory Flash condition. Second, this illusion was less strong, being successful on 77% of trials (vs. 83% of illusory flash trials; χ^2^(2391) = 24.83, p < .001). In fact, one participant had to be rejected from this analysis due to an insufficient number of Suppressed Flash illusions (see Methods). Despite this, we found that activity patterns in the deep layers of V1 were more similar to those on trials in which no middle flash was presented (i.e., two flashes and two beeps, similar perceptual outcome) than to those in which a middle flash was presented (i.e., three flashes and three beeps, same retinal input; one-sided t-test: t(21) = 1.926, *p* = .034; Figure 4A). This effect was, again, not present in deep layer voxels tuned to a control location (all layers *p* > .05; 4B). The middle and superficial layers did not show the same effect (middle: t(21) = .516, *p* = .695; superficial: t(21) = −0.568, *p* = .288), nor did middle layers differ significantly from superficial or deep layers (deep vs. middle: t(21) = −1.682, *p* = .107, superficial vs. middle: t(21) = −0.945, *p* = .356).

**Figure 4.**
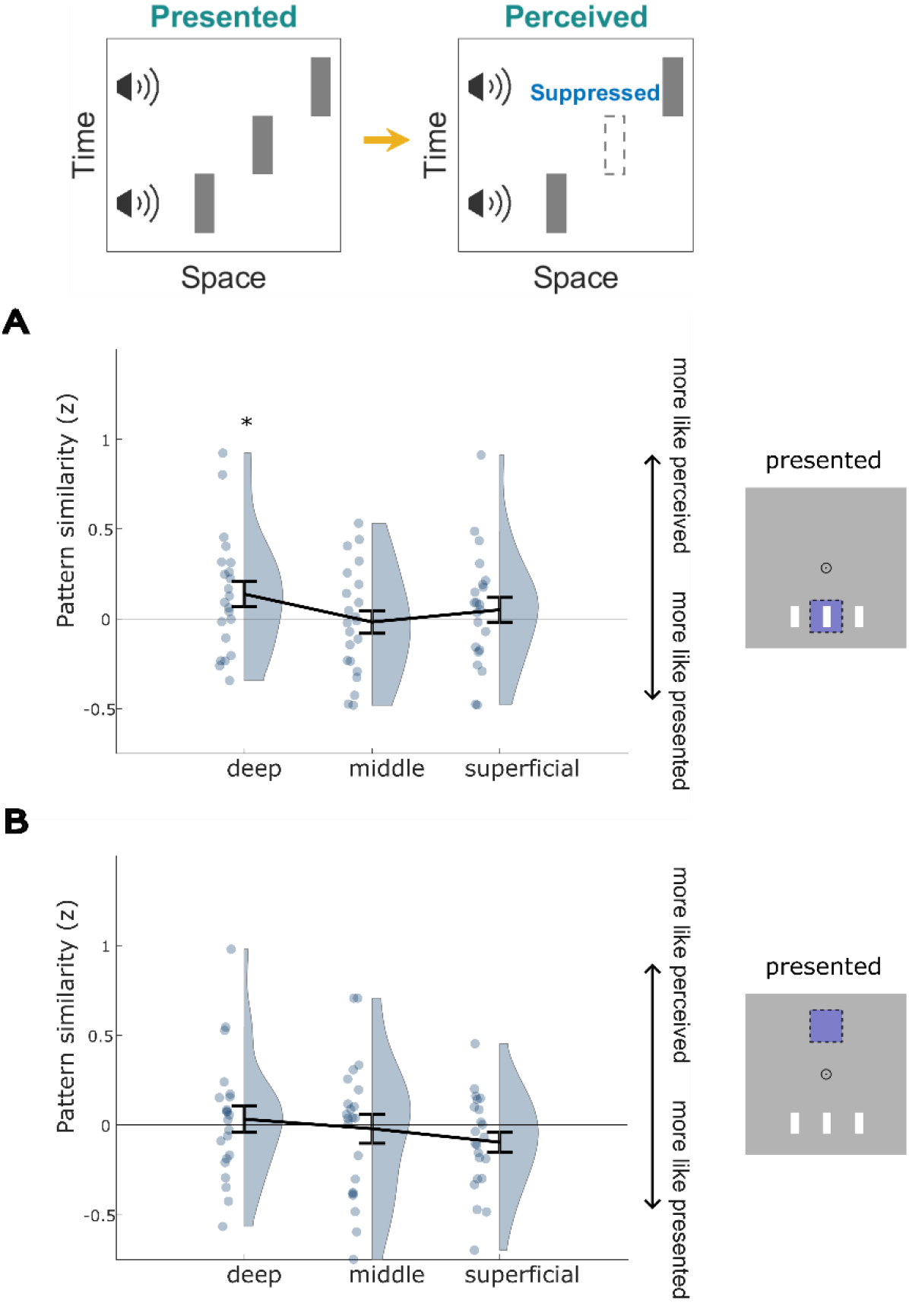
Perceptual pattern similarity in BOLD activity patterns for voxels tuned to the second flash and control location per layer of V1 for the ‘Suppressed Flash’ illusion (left); the ROI is indicated on screen relative to perceived stimuli (right). A) Perceptual pattern similarity of voxels tuned to the illusory flash location. Positive values indicate that activity patterns were more similar to activity patterns evoked by stimuli inducing the same percept (‘No Flash’), while negative values indicate more similarity to conditions with identical retinal stimulus but different percepts (‘Real Flash’). B) The same analysis for voxels tuned to a control retinotopic location. Error bars denote SEM across participants. * *p* < 0.05.

Are these pattern similarity effects caused by illusory percepts activating the same area of cortex as veridical percepts do? To test this, we probed whether the overall BOLD amplitude (rather than activity patterns) was affected by illusory percepts (S3). First, we tested whether the ‘Illusory Flash’ condition upregulated BOLD activity compared to the control condition ‘No Flash’ – again, only for voxels whose receptive field covers the middle flash location. We did not find evidence for this (‘Illusory Flash’ > ‘No Flash’: t(22) = −.316, *p* = .0.623; Figure S3A), while a real flash did increase BOLD activity (‘Real Flash’ > ‘No Flash’; t(22) = 2.184, *p* = .02) – validating our approach. Conversely, if perception is reflected in the overall BOLD amplitude, the ‘Suppressed Flash’ condition should evoke a lower BOLD amplitude than a ‘Real Flash’,just like the ‘No Flash’ condition. We found no evidence for a perception-driven amplitude change here either (‘Suppressed Flash’ < ‘Real Flash’: t(21) = −1.306, *p* = .897). There was also no evidence for BOLD up- or downregulation within the deep layers specifically, where the pattern similarity effects were present (S3B; ‘Illusory Flash’ > ‘No Flash’: t(22) = −1.076, *p* = .853; ‘Suppressed Flash’ < ‘Real Flash’: t(21) = −.907, *p* = .187; Figure S3). Taken together, the findings indicate that subjective perception was reflected in activity patterns, but not in the overall amplitude of the signal.

How can we reconcile these pattern similarity effects in the deep layers with the absence of an overall BOLD amplitude effect (Figure 3D and 4D)? Focussing on the ‘Illusory Flash’ condition, one potential explanation could be that the illusory flash leads to an increased neural signal at voxels tuned to the exact illusory flash location, flanked by suppression on either side (e.g., due to lateral inhibition). This could result in a net zero signal in the region between the two real flashes, while retaining retinotopically precise illusion-induced activity. If this were true, one would expect to observe a negative correlation between illusion-induced BOLD amplitude and the distance of a voxel’s receptive field centre from the illusory flash location in V1 deep layers. We tested this post-hoc hypothesis by calculating the correlation between BOLD amplitude and absolute horizontal distance between the centre of the middle flash location and voxels’ pRF centre. Indeed, we found a negative correlation for the Illusory Flash condition (r = −.11) but not the No Flash condition (r = −.01) in the deep layers of V1 (Illusory Flash vs. No Flash; t(22) = 1.840, *p* = .039). While post-hoc, these results are in line with the illusion inducing highly retinotopically specific neural activity, flanked by suppression [15,23].

Which brain area might be the source of this feedback-induced activity in V1? The superior temporal cortex (STC) is a prime candidate, given its role in multisensory integration [24–27] and suggested role in postdiction [4]. We additionally investigated the hippocampus as a more exploratory target, given its role in representing predictive sequences [28,29]. We calculated informational connectivity to establish whether perceptual information was shared between areas across trials. Strikingly, we found that for the ‘Illusory Flash’ condition, perceptual pattern similarity in superior temporal cortex covaried more with V1 deep layer voxels than with middle layer voxels tuned to the location of the flash sequence, i.e. the three flash locations (two-sided t-test; t(22) = 2.562, *p* = .018; Figure 5A). This did not apply to a control retinotopic location (two-sided t-test; t(22) = −1.251, *p* = .225; Figure 5B). This effect was not significantly present for superficial layers (‘flash 123’, superficial vs. middle layers: t(22) = 1.659, *p* = .112; control: t(22) = −0.654, *p* = .519). This finding is in line with the hypothesis that STC communicates information about post-hoc inferred illusory flashes back to V1. Note that this effect was not present for the ‘Suppressed Flash’ illusion (‘flash 123’: *p* > .05 for deep vs. middle and superficial vs. middle; control: *p* > .05 for deep vs. middle and superficial vs. middle), possibly due to a combination of lower illusion susceptibility and the fact that top-down and bottom-up signals are less cleanly separated in this condition (see above). There was also no layer-specific informational connectivity between V1 and the hippocampus during the illusion (‘flash 123’ deep vs. middle: t(21) = 0.566, *p* = .578; ‘flash 123’ superficial vs. middle: t(21) = 1.265, *p* = .220).

**Figure 5.**
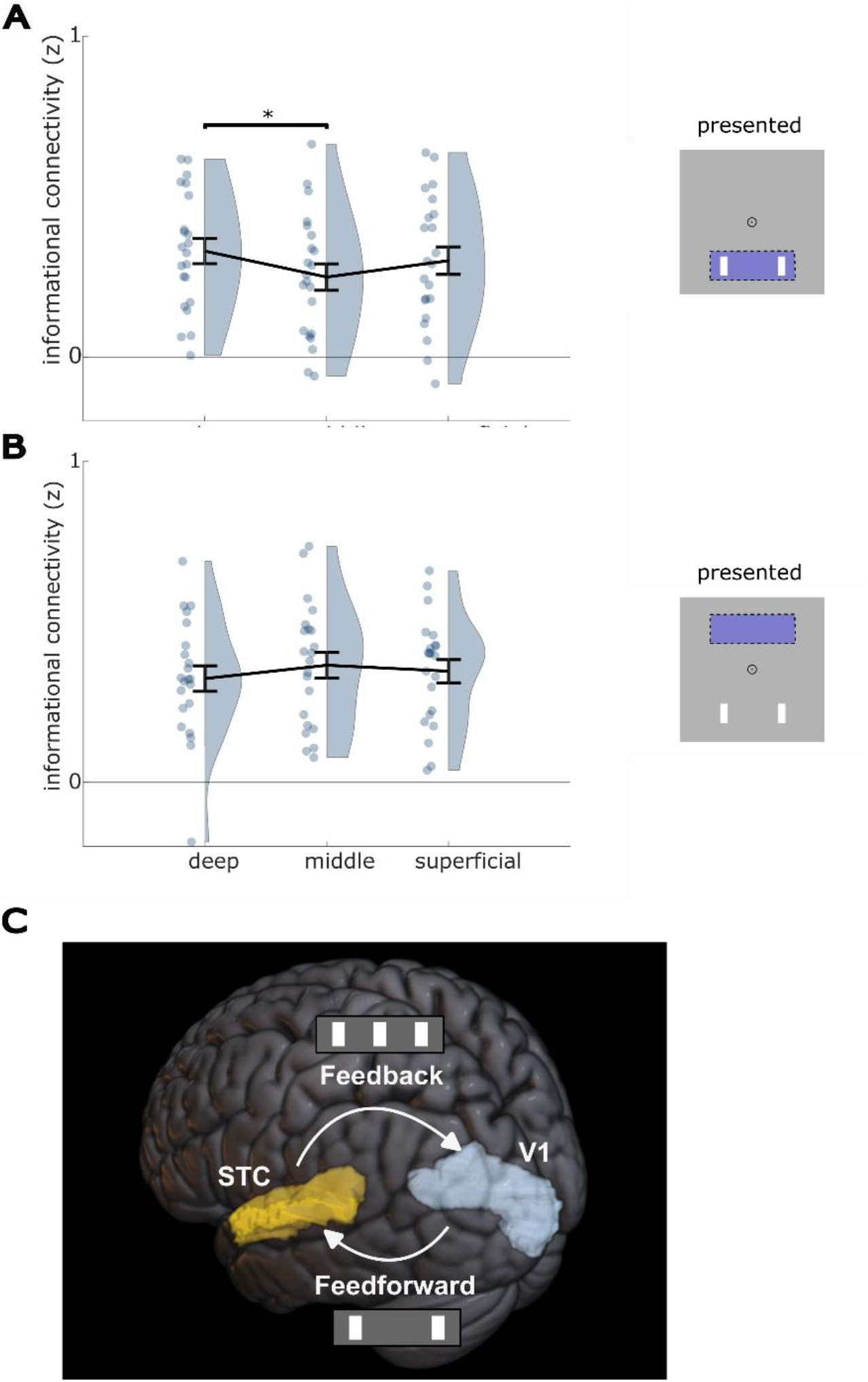
Informational connectivity (IC) between STC and V1 layers for the ‘Illusory Flash’ illusion. A) IC between STC and V1 voxels tuned to the flash sequence (union of left, middle and right flash locations) on the left. Higher values denote more shared perceptual pattern similarity across trials. Error bars denote SEM across participants. * *p* < 0.05. B) the same analysis for voxels tuned to a control location. C) A schematic figure of how V1 at first processes presented visual stimuli, but may later receive an updated percept from STC.

## Discussion

Here, we investigated the neural mechanisms of postdiction – a phenomenon in which the perception of a stimulus is shaped by later information. We found that postdictive illusions induced activity patterns in the deep layers of V1 that resembled those induced by veridical perception (as opposed to equivalent retinal input). Strikingly, this was true for two opposite postdictive effects: inducing and suppressing flashes – providing support for the generality of this neural mechanism. The postdictive effects were reflected only in voxels tuned to the illusory flash location, demonstrating a retinotopically specific neural correlate of the illusory percepts. The effect was not present in the middle layers of V1, in line with the fact that middle layers predominantly receive bottom-up retinal input and are avoided by top-down feedback [22].

The effect was also not significantly present in superficial layers, which do receive feedback inputs [22]. Apart from that the signal may be diluted by feedforward inputs as discussed above, it may be that the type of feedback studied here specifically targets deep cortical layers because it arrives from relatively distant sources [30], as has been reported for other feedback signals like prediction [19] (but see Muckli et al., 2015 [31]), surface filling-in [32], and mental imagery [33]. Alternatively, it has been suggested that deep layer neurons represent perceptual hypotheses (here, inferred visual flashes), while superficial layer neurons respond to mismatches between those hypotheses and inputs (here, difference between inferred and presented flashes), i.e., prediction errors [34–36].

The presentation of visual stimuli typically evokes a peak in neural activity in sensory cortex, as confirmed for the real flashes presented in the current study. However, for the illusory percepts studied here there was a close link between perception and the *pattern* of neural activity, but not its overall amplitude. This finding is in line with intracranial EEG research showing that sustained perception is reflected in neural activity patterns rather than in response amplitude [37]. It should be noted that a post-hoc analysis revealed that the illusory flash-induced activity pattern in V1 may in fact reflect a highly spatially specific amplitude increase, but encoded more sparsely than a retinally presented flash and possibly flanked by inhibition [38]. Taken together, the evidence across illusions, layers and retinotopic locations is strongly in favour of a feedback model of perceptual temporal integration, where deep layers of primary sensory cortex receive postdictively inferred perceptual information through top-down connections.

These findings diverge strongly from the traditional view of V1 as a passive feature extractor that processes sensory information coming in through the retina and sends it onwards to higher-order visual regions [39,40]. Plenty of evidence over the last few decades has demonstrated that activity in V1 can be modulated by top-down processes like attention and expectation [41–43]. However, this work has focused almost exclusively on how preparatory top-down processes modulate early neural responses (but see Sergent et al., 2013 [44]). Our findings demonstrating a role for postdictive feedback to V1 suggest that it is still involved in perceptual inference at a late stage of sensory processing, far beyond the initial feedforward sweep. This role suggests that early visual cortex closely tracks visual perceptual experience, even in the absence of retinal stimulation.

One open question is whether V1 also reflects the temporal order of perception. Although the illusory flash is only inferred after all the real flashes (and tones) have been processed, it is perceived as occurring simultaneous with the second tone, and prior to the third tone-flash pair. This results in a distinction between time of processing and time of perceiving [3]. Given the lack of temporal resolution of fMRI, we currently cannot say which of these timelines is reflected in V1. Future research using time-resolved methods such as EEG/MEG should test whether activity in V1 reflects time of perceiving or time of processing – or both.

We hypothesized that multisensory areas higher up the cortical hierarchy integrate audiovisual signals and feed information resulting from post-hoc perceptual inference back to V1 [4], similarly to predictive coding [35,45]. In line with this, we found that perceptual pattern similarity in STC and V1 deep layers co-fluctuated, implying that STC indeed may send postdictively inferred information to V1, revising early sensory representations. STC is well-positioned to fulfil this function given its role in multisensory integration [46]. Along with involvement in audio-visual predictions [24,25], it suggests a role in evaluating and updating sensory input. Importantly for postdiction, STC not only has large spatial receptive fields, but also large temporal receptive fields [12]. Together, these properties allow STC to track the rich spatiotemporal information necessary for postdictive inference, in line with previous suggestions [4]. Recent work suggests that the illusions used in this study are best explained by a postdictive causal inference model [47]. Given the neural properties discussed above, STC may fulfil this function. This question, however, remains open for future research.

### Conclusion

Postdictive effects demonstrate that perception is not a livestream of reality, but rather a post-hoc montage. Here, we demonstrate that this montage affects neural representations in the primary visual cortex, as a result of feedback from higher-order multisensory cortex. Thereby, the current study sheds light on how our subjective experience of the world is constructed through perceptual inference across space and time.

## Methods

### Participants

Seventy-five human volunteers of both sexes with normal vision and hearing gave written informed consent and participated in the study. Participants were excluded after the behavioural screening if they scored below 85% correct on control trials and if they were less than 40% susceptible to the induced illusion. Twenty-four participants were scanned during the neuroimaging session. After neuroimaging, participants were also excluded if they failed to meet strict head motion criteria of maximum 10 movements of 1 mm or greater between successive functional volumes. If participants did not have more than ten successful illusions across scanning runs for a given illusion, the participant was excluded for analysis of that illusion. Twenty-three participants (four males, mean age 23.6 ± 4.0 years) were included using these criteria for analysis of the ‘Illusory Flash’ illusion, with one exclusion due to head motion. Twenty-two participants were included for analysis of the ‘Suppressed Flash’ illusion, since one extra participant failed to meet the illusory susceptibility threshold for this illusion in the scanner. Participants received £12 per hour monetary compensation for their time. The study was approved by the Research Ethics Committee at UCL.

### Stimuli

Stimuli were generated using MATLAB 2019a [48] (MathWorks, Natick, MA, USA) and Psychophysics Toolbox Version 3 [49].

In the behavioural screening, participants were sat in front of a computer, where stimuli were displayed on aLet Dell Optiplex 3070. Operating System: Windows 10 64-bit. Monitor: Dell UltraSharp U2415 with a resolution of 1920 × 1200 and a refresh rate of 60Hz. Sound was presented from two speakers from each side of the computer.

In the MRI scanner, visual stimuli were displayed on a rear projection screen using an Epson EB-L1100U projector (1920 × 1200 resolution, 355 x 220 mm, 60-Hz refresh rate) against a gray background. Participants viewed the visual display through a mirror that was mounted on the head coil with a viewing distance of 113cm, with a resulting visual field of 17.9° horizontal and 11.1° vertical. Etymotic Ear-Tone ER3A systems have been adapted to present auditory stimuli in MRI environments. We adjusted the volume of the auditory stimuli so that participants could clearly hear them over the scanner noise, without being uncomfortable loud. We verified audio-visual stimulus presentation timing using an oscilloscope.

Stimuli were adapted from [5].The visual stimuli were white rectangles (or ‘flashes’) on a dark grey background, presented 4.3° below a bull’s eye-shaped fixation point. The flashes subtended 0.28° × 1.2° visual angle (width x length) each. Horizontal distance between flash centres was 2.13° and flashes were shown one-by-one from left to right for a duration of 16.6ms each with 52ms in between flashes. The auditory stimuli were 800 Hz tones presented for 10ms each, modulated by a square wave. The auditory tones were presented synchronously with the visual flashes, accounting for the presentation latency of the projector used in the MRI scanner. Before stimulus presentation, a 100ms pre-stimulus window alerted the participant by turning the central dot of the fixation bull’s eye from light grey to dark grey. After stimulus presentation, participants had 2000ms on each trial to report whether they saw two or three flashes, and how confident they were in their answer (4: very confident, 3: quite confident, 2: not so confident, 1: not confident at all). No visual response cues were provided, but instead they were instructed and trained beforehand to respond in this manner. If one of the responses was missing, the central dot of the bull’s eye briefly flickered red. Finally, the central dot turned light grey to indicate the trial had ended. A jittered inter-trial-interval (ITI) of 1600ms, 3100ms or 4600ms (respectively, 50%, 30% and 20% chance) preceded the next trial.

### Experimental design

We used the Audiovisual Rabbit paradigm [5] to induce two complementary audio-to-visual illusions by pairing the flashes with the sounds (Figure 1). The illusions have in common that, when successful, the presence or absence of the second beep dictates what is seen. [5] demonstrated the postdictive nature of these illusions; e.g. the illusion only occurs if the third beep is presented, while the illusory flash is perceived simultaneous to the second beep, and the location of the third flash determines the location of the illusory second flash. These illusions provide the opportunity to isolate neural activity induced by the illusion from any retinal information, especially the illusory flash, which was perceived in absence of any corresponding retinal input.

We added two control conditions, to orthogonalise sensory input and perception in a 2-by-2 design (Figure S1). In one condition both the middle flash and middle tone were omitted, presenting only two flash-beep pairs (‘No Flash’), whereas in the other control condition three flash-keep pairs were presented (‘Real Flash’).

### Procedure

Participants first completed a behavioural practice and screening session on day one. Here, participants read instructions off the computer screen that explained the objective of the experiment without giving away the illusions: to focus on the flashes while fixating on the centre of the screen. All four conditions were presented equally often, interleaved in pseudorandomized order. The participants completed two blocks of 32 practice trials, receiving verbal feedback after each block in the rare cases where they frequently provided no answers or answered too late. Next, they completed four runs of 52 trials for a total of 26 per condition. Those susceptible to the illusion (see criteria in ‘Participants’) were invited for the neuroimaging session.

During the fMRI session the same experiment was presented for four run of 104 trials each. Additionally, we conducted four runs of pRF mapping. Here, participants fixated on a central fixation dot while bars of black-and-white checkerboard stimuli travelled across the screen within a 5.5 degree radius from fixation [21]. Specifically, we used the ‘8 bars’ stimulus set in MrVista, which were presented in four runs of 5.5 minutes each.

### Behavioural analysis

We tested for successful visual illusions by comparing them to the condition with the same retinal stimulation, but differing in sound – which induces the difference in perception. For the ‘Illusory Flash’ condition, we compared whether more flashes were reported than in the ‘No Flash’ condition, and for the ‘Suppressed Flash’ illusion, we asked whether fewer flashes were reported than in the ‘Real Flash’ condition. Both analyses were done with two-sided Wilcoxon rank sums test for non-normally distributed data. We also quantified participants’ level of confidence in their percepts, both real and illusory.

### Scanner protocol & sequences

MRI images were acquired on a 7T MAGNETOM Terra (Siemens Healthcare, Erlangen, Germany) scanner (with a head coil equipped with 8 transmit and 32 receiver channels (Nova Medical, Wilmington, USA) at the Functional Imaging Laboratory (University College London). Partial-brain functional images were collected with a 3D EPI protocol (T2* weighted, Volume acquisition time = 3.264 seconds, TR = 68.00ms, TE = 26.08, Spatial resolution: 0.8 mm isotropic. Field of view: Angled to optimize coverage of visual cortex, temporal lobe, and hippocampus. Angle adjusted individually for each participant). Four EPI runs were collected with variable volumes due to intertrial jitter (~190-210 volumes), and 113 volumes per run for pRF mapping. Four dummy volumes with the opposite PE direction were acquired at the start of run, these were not included in the preprocessing pipeline.

We collected structural data through Magnetization Prepared 2 Rapid Acquisition Gradient Echoes (MP2RAGE) with a TR of 68ms and TE of 2.54ms, with a modified slab size of 288 slices instead of 240 to account for larger heads – angled at 15.0 degrees. Fat suppression was applied for artefact prevention.

### Neuroimaging preprocessing

#### MP2RAGE

Structural brain scans were preprocessed with FreeSurfer version 7.3.2 [50]. The brain image was skull stripped and dura was automatically removed with ‘synth strip’, a deep learning-based algorithm [51]. Removal of remaining dura was done manually before approximating the pial and white matter layers to aid the process. Using automatic parcellation provided by Freesurfer, we extracted anatomical labels for primary visual cortex, hippocampus and superior temporal cortex, that were used to extract voxels for our analysis.

#### Cortical layer definition

We subdivided cortex into three equivolumous layers using the level set method [52] supported by the idea that cortical layers maintain volume throughout the gyri and sulci. The cortical volume was used to compute the distance between white matter and pial surfaces, taking the local curvature into account. For each voxel we then computed to what extent it was located within each of five compartments (deep, middle and superficial gray matter layers, white matter, and cerebrospinal fluid). Each voxel was assigned to one of the three gray matter layers if >50% of its volume resided within that layer.

#### fMRI preprocessing

The 3D EPI data were preprocessed using Statistical Parametric Mapping [53] version 12 (SPM12). NORDIC was applied to remove thermal noise and improve SnR [54]. Realignment was done to estimate - and correct for - motion. We applied no spatial smoothing to preserve high spatial resolution. The structural data were coregistered to the mean EPI volume using a two-step process. First, volume-based coregistration in SPM12 was applied, and subsequently a recursive boundary-based registration [55] (RBR) was used to correct for distortions in the phase encoding direction. RBR first applies an affine boundary-based registration (BBR) recursively to increasingly smaller partitions of cortex with seven degrees of freedom, rotation and translation in the three dimensions, only scaling along the phase-encoding direction to allow distortion correction. In each of six recursions, the currently selected cortical mesh was split into two and the optimal registration was applied to both parts, before each part was subdivided into two and registered again; increasing the specificity and the fit between structural and functional volume (that are distorted by magnetic field inhomogeneity) at each stage. The splits were made along the cardinal axes to obtain equal numbers of vertices for each part.

From the pRF fMRI data, we used mrVista [56] to estimate for each voxel a population receptive field in the form of a two-dimensional Gaussian distribution with a centre (x and y coordinate) and a standard deviation (sigma). We excluded voxels that had low variance explained (r^2^ < 10%) during the pRF task, had an unrealistically small receptive field (sigma < .2 visual degree angle) or too large a receptive field to have the spatial specificity our analysis demanded (sigma ≥ 1.3 visual degree angle).

Following pRF estimation, we visually checked whether the estimated maps corresponded to the expected retinotopic organization of visual cortex by inspecting maps of x-coordinates, y-coordinates and receptive field sizes. The pRF estimates were used to select voxels tuned to particular locations in the visual field.

#### Retinotopic locations and ROI selection

We selected V1 voxels whose receptive field fell on specific regions of interest in the visual field (S2). The primary retinotopic location of interest was the location of the middle flash, i.e. the location where the illusory flash was perceived (on Illusory Flash trials). As a control location, we used the mirror image flipped over the y-axis, so that the selected voxels for each location had similar neural properties due to the equal distance from the fixation point. Per location, we selected voxels whose centre of receptive field (x,y) was within circumference *sigma* of the flash location. Debriefing after the fMRI session (specifically, answering: ‘did you see the second flash exactly in the middle?’) revealed individual variability in reported location of the illusory flash. Specifically, of 23 included participants, three reported seeing the middle flash exactly in the middle at all times, whereas the other twenty participants reported seeing it at least sometimes to the left and/or right of centre as well. Therefore, the ‘flash 2’ region of interest was broadened four-fold horizontally to include as many voxels as possible whose receptive field covered the illusory percept, while avoiding any overlap with the other two flashes. Finally, this retinotopic ROI was subdivided into three cortical layers, – deep, middle, superficial, see above – with respective voxel counts (mean ± SEM) of 89.30 ± 14.25, 77.61 ± 12.70, 87.22 ± 17.65.

#### First level analysis

For the task fMRI data, we applied a General Linear Model (GLM) using SPM12 to model condition-specific BOLD responses for each run. In this GLM we only included control trials (‘No Flash’ and ‘Real Flash’) on which participants responded correctly, and illusion trials (‘Illusory Flash’ and ‘Suppressed Flash’) on which participants experienced the illusion. Incorrect and missed responses were modelled with a separate regressor, whose parameter estimates were not analysed. The head motion parameters, their derivatives, and the square of the derivatives were included as nuisance regressors. Subsequently, the data and the design matrix were high-pass filtered (cut-off = 128 s) to remove any low-frequency signal drifts. For each participant and each condition, the resulting estimated beta parameters per voxel were averaged across runs, resulting in four sets of parameter estimates per individual. The retinotopic ROIs described above were used to extract voxel activity patterns for each condition and each ROI, which were subdivided into three cortical layers.

#### Multivariate analysis

To analyse whether illusions altered neural activity in primary visual cortex, we implemented both a multivariate and a univariate analysis. In the multivariate analysis, we computed a ‘perceptual pattern similarity’ metric: the pattern similarity between the activity during illusory trials and the control condition with the perceptual equivalent minus the similarity with that of the retinal equivalent (Figure S1).

This involved calculating the Pearson correlation between estimated BOLD activity patterns for different conditions for V1 voxels tuned to the middle flash location – i.e. the location where illusions sometimes occured. We used Fisher’s z transformation to create normally distributed correlation scores. For the ‘Illusory Flash’ illusion, for example, we calculated per participant whether the pattern of brain activity when perceiving an illusion was more similar to that when viewing a real flash than to when no flash is presented. That is, we subtracted the correlation between ‘Illusory Flash’ and ‘No Flash’ from the correlation between ‘Illusory Flash’ and ‘Real Flash’. To analyse the (differential) impact of postdictive perception on V1 layers, we performed a two-way repeated-measures ANOVA, with both ‘Layer’ (levels: deep, middle, superficial) and ‘Illusion type’ (levels: ‘Illusory Flash’, ‘Suppressed Flash’) as within-subject factors. We followed up with two-sided t-tests to analyse how perceptual information averaged over illusions differed between layers. Lastly, we evaluated our hypothesis that deep layers encoded this postdictive perceptual information by testing whether it was significantly higher than zero, using a one-sided student’s t-test. We applied the same analysis to the retinotopic control location for validation, and evaluated the same hypothesis for each illusion individually.

#### Univariate analysis

In the univariate analysis, we compared the amplitude of BOLD responses of voxels tuned to the middle flash location across the ‘Illusory Flash’ condition and ‘No Flash’ condition by averaging them per participant and applying a one-sided student’s t-test across participants. For validation we compared the middle flash location to its mirror image above fixation during the ‘Real Flash’ condition.

#### Informational connectivity

To investigate the source of the illusory signals, we implemented an informational connectivity analysis [57,58] to probe trial-by-trial covariation in pattern similarity (correlation with ‘Real Flash’ minus correlation with ‘No Flash’) between two areas. To optimise the robustness of the analysis, we enlarged the retinotopic location of interest to include all three flashes and thus the signal-to-noise ratio, increasing the voxel count per layer (mean ± SEM): deep (179.04 ± 20.22); middle (160.22 ± 19.49); superficial (192.00 ± 26.31). To obtain a perceptual information metric per trial, we computed the mean pattern for the ‘No Flash’ and the ‘Real Flash’ control conditions separately per participant, and calculated on a trial-by-trial basis how similar the neural responses during the illusory conditions were to one control condition versus the other, to measure perceptual information. We extracted this pattern per ROI and correlated the across-trial patterns between pairs of ROIs. This resulted in connectivity measures between brain areas for each participant, that were normalised using Fisher’s z transformation. To analyse which area might feed back perceptual information to primary visual cortex, we analysed informational connectivity per V1 layer. We compared the perceptual information shared between each of candidate areas superior temporal cortex (voxels: 28,787 ± 645.60), and hippocampus (voxels: 12,638 ± 179.86), and each V1 layer using one-sided student’s t-tests. We performed the same analysis for the retinotopic control location – a mirror image of the three flashes – for validation.

## Notes

### Competing Interest Statement

The authors have declared no competing interest.

### Summary of Updates

Updated ORCID ID of co-author; no other changes were made.

