## supplementary_figures for "The post-hoc montage of perception: deep layers of primary visual cortex encode postdictive percepts"

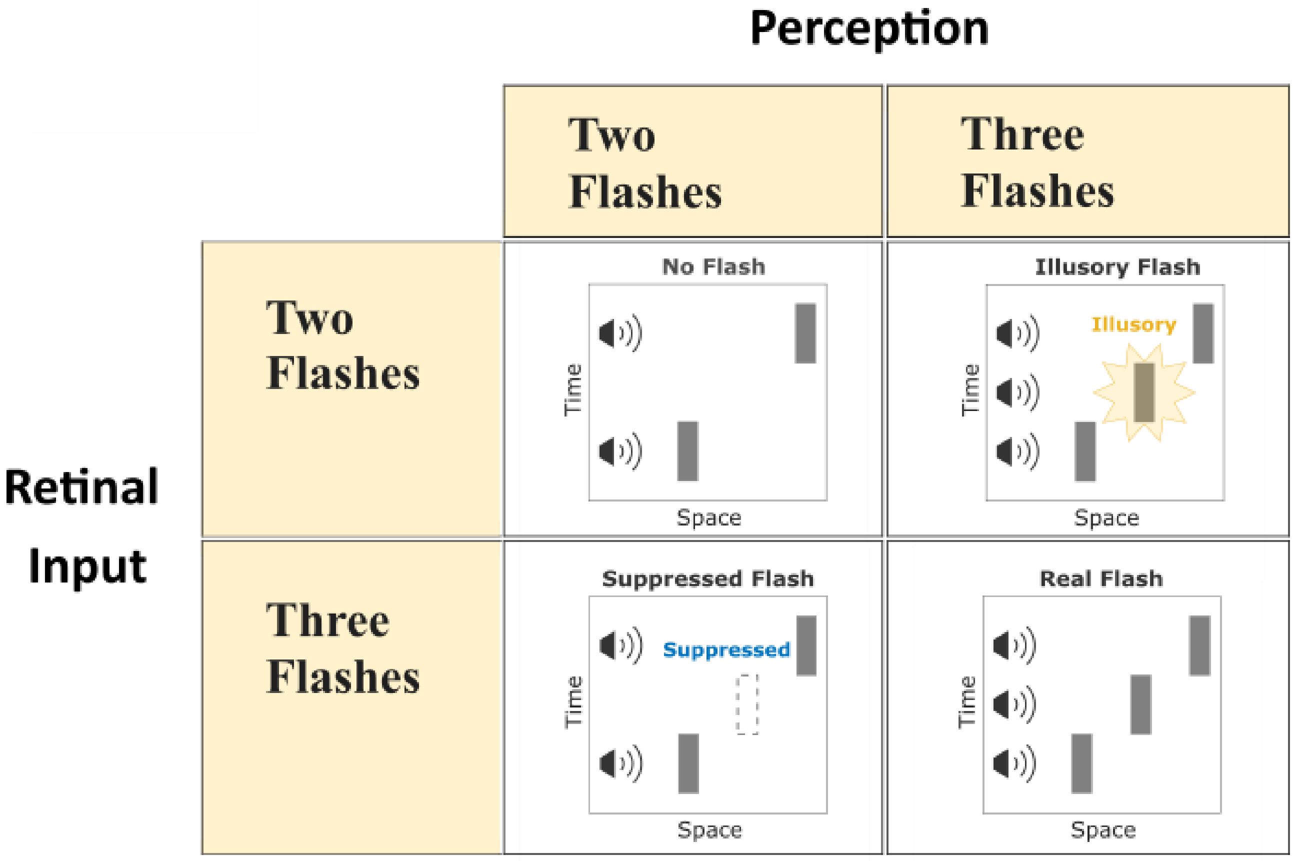


*Figure S1.* The experiment contained four conditions, forming a 2x2 design across retinal input and perception. This design allowed comparison of illusions to conditions with the same perception but different retinal input and contrast that against comparison with conditions with different perception but identical retinal input.

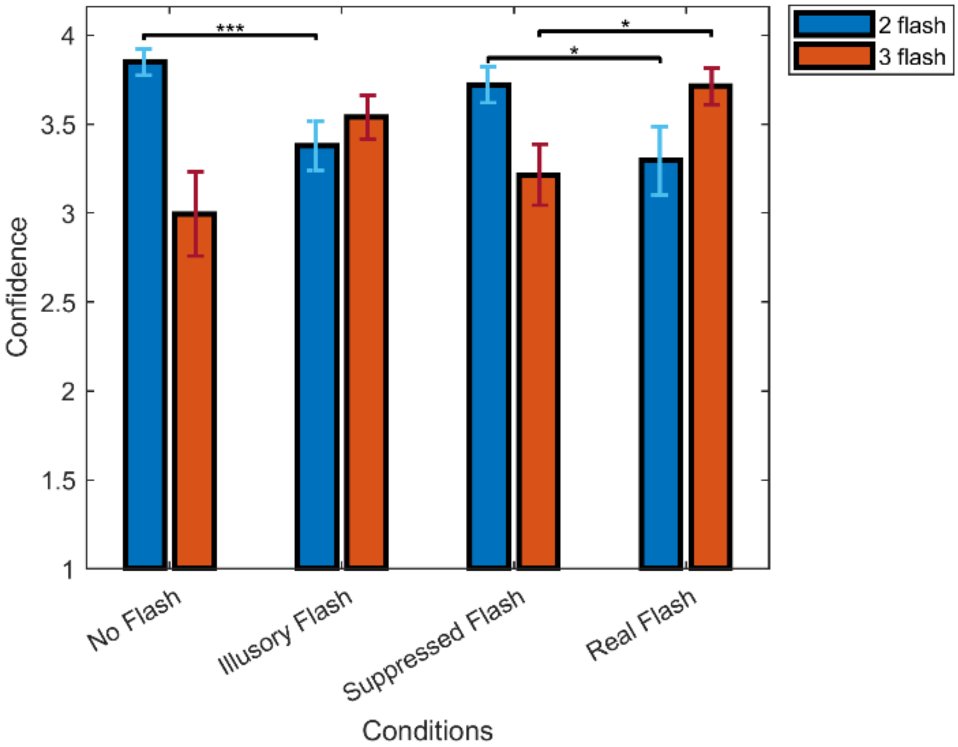


*Figure S2.* Confidence is plotted for trials with two reported flashes and for three reported flashes per condition. We compared both illusions with the control condition with the same retinal input, but different auditory input – and consequently, potential differences in perception. We found that for the 'Illusory Flash' an additional auditory stimulus decreased the confidence in two flash reports compared to 'No Flash' as tested with a two-sided t-test (t(23) = -4.460, $p < .001$), while it did not show confidence increase in three flash reports (t(23) = 1.767, $p = .094$) – most likely due to lack of three flash reports in the 'No Flash' condition. Conversely, for the 'Suppressed Flash', an omitted auditory stimulus decreased the confidence in three flash reports compared to 'Real Flash' (t(23) = -2.545, $p = .020$), and increased the confidence in two flash reports (t(23) = 2.592, $p = .017$). Bars denote SEM across participants. * $p < 0.05$; *** $p < 0.001$.

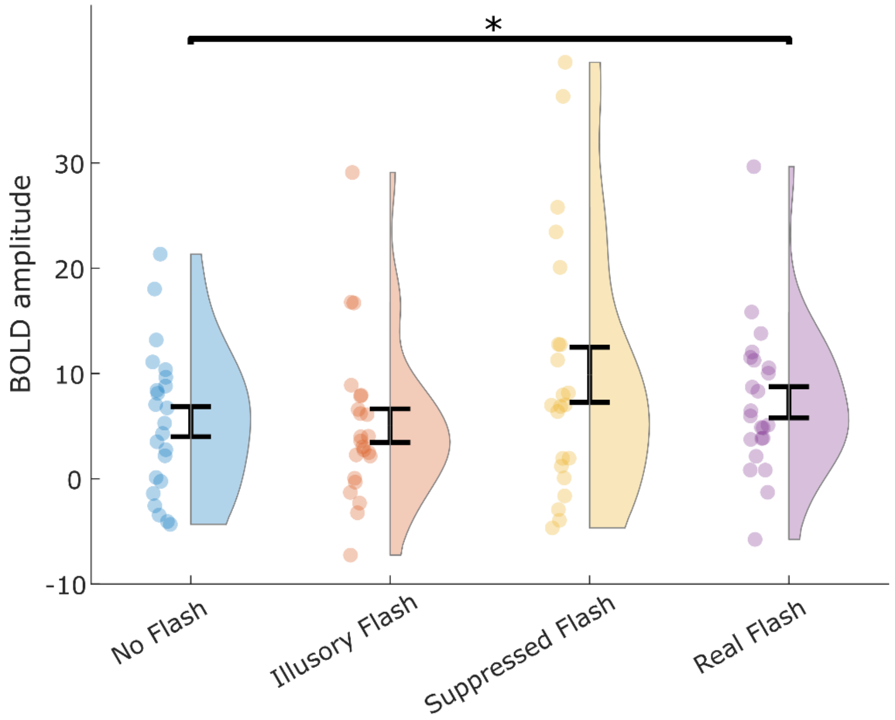


*Figure S3.* The BOLD amplitude for voxels in V1 and V1 deep layers per condition tuned to the 'Flash 2' location. A) All of V1: Perceiving an illusory flash did not upregulate BOLD amplitude compared to perceiving no flash ('No Flash' condition), and perceiving a suppressed flash did not downregulate BOLD amplitude compared to perceiving that flash ('Real Flash' condition), despite that a real flash did upregulate BOLD amplitude compared to no flash being presented ('Real Flash' > 'No Flash'). B) V1 deep layers: illusory perception again did not change overall BOLD amplitude, despite matched retinal input and evidence of BOLD modulation in deep layers for both illusions. * $p < 0.05$.
